## Supplemental Figures and Legends for "Hierarchical membrane-chromatin tethering buffers nuclear envelope assembly against alterations in lipid flux"

**Supplemental Figures index**

Supplemental Figures and Figure legends S1-S7

Supplemental Movie Legends Movie S1-S9

Tables S1-3

#### Supplementary Figures

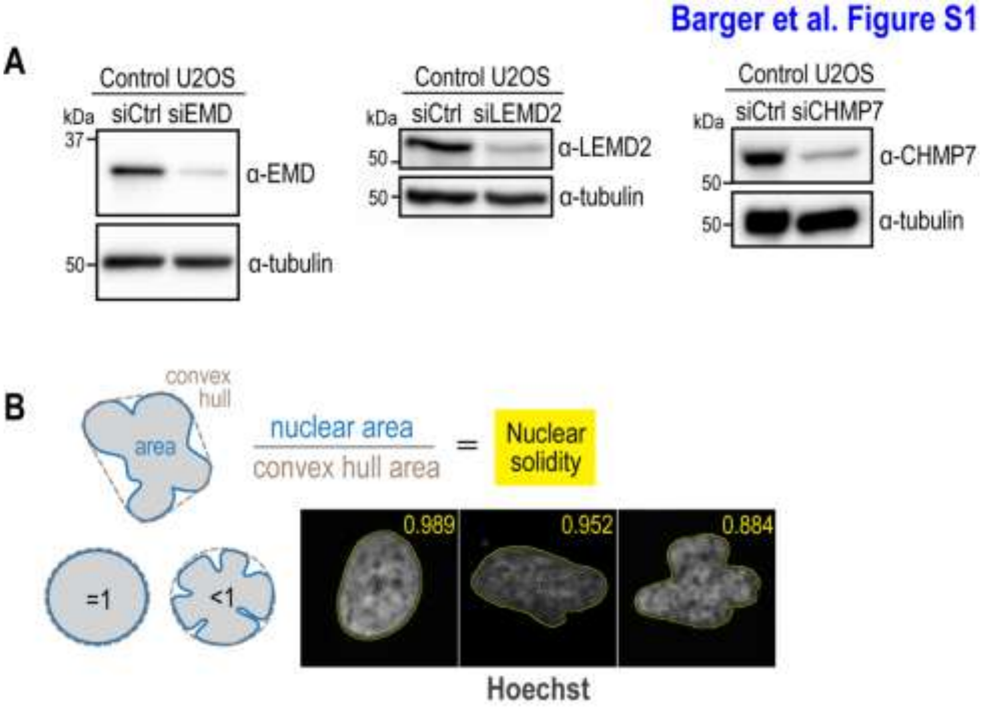


**Figure S1.** (A) Representative immunoblots of siRNA knockdowns for Emerin, LEMD2, and CHMP7 in Control U2OS cells. (B) Schematic of nuclear solidity quantification and example images.

**Barger et al. Figure S2**

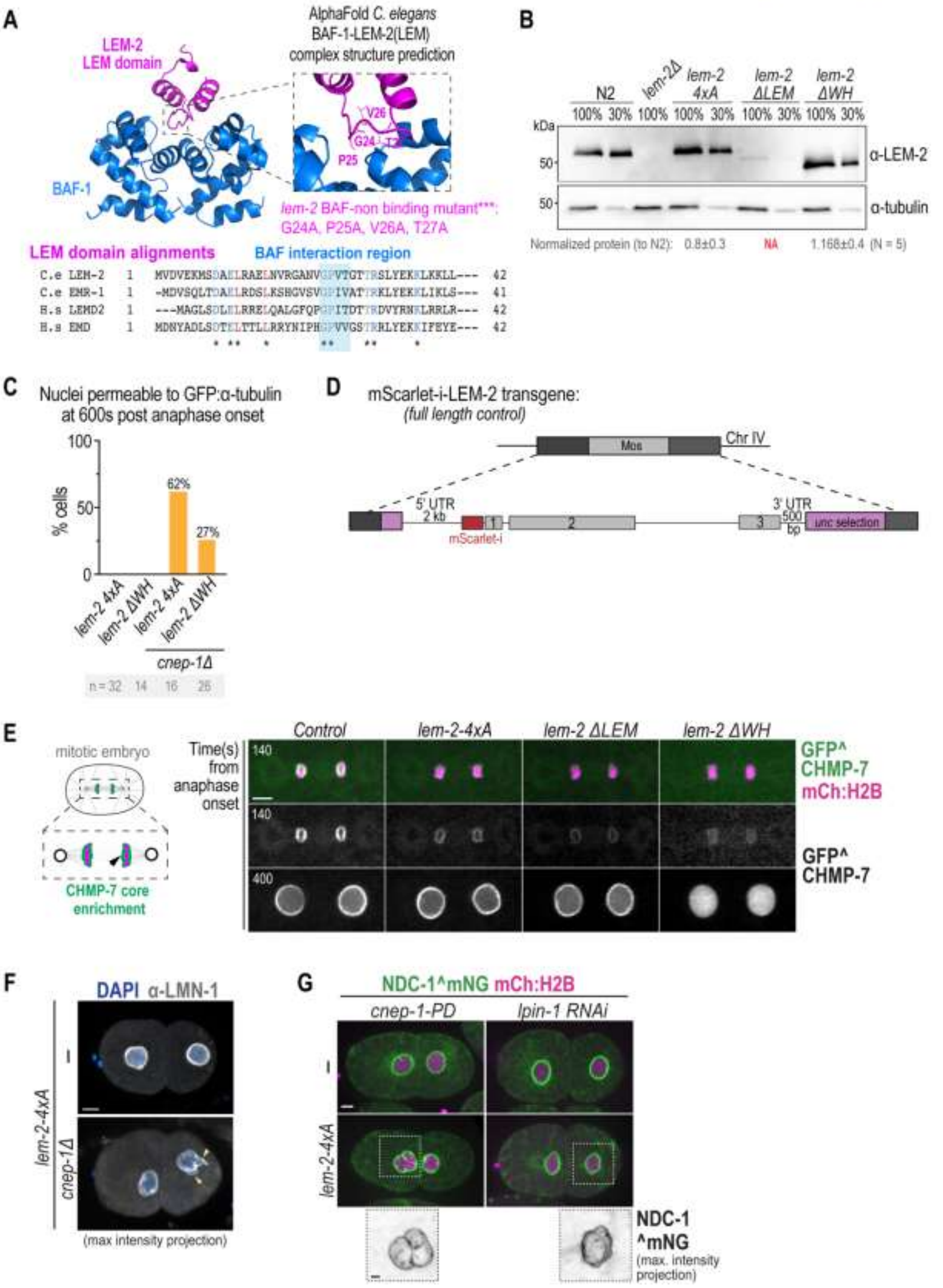

**Figure S2.** (A) *C. elegans* BAF-1 homodimer (blue) with LEM-2 LEM domain (pink) complex structure prediction by AlphaFold 3 (Abramson *et al.*, 2024). Zoom in shows position of 4 amino acid substitution residues for BAF-nonbinding mutant. Below, amino acid sequence alignment of LEM domain among indicated proteins. *indicates identical amino acid residues. Blue shading marks BAF-interacting region mutated in the *lem-2-4xA* and *emr-1-4xA*. (B) Representative immunoblot of whole worm lysates incubated with indicated antibodies in indicated strains. Below, quantification of LEM-2 expression by immunoblot normalized to N2 levels (N = 5 biological replicates). To note, LEM-2 antibody is made against aa1-100 including the LEM-domain and is unable to fully recognize the *lem-2 ΔLEM* strain. (C) Plot representing percentage of nuclei permeable to GFP:α-tubulin at 600 sec post anaphase onset (n = # of cells). (D) Schematic showing single copy insertion of mScarlet-i-tagged LEM-2 transgene integrated in the MosI transposon site of chromosome IV, later crossed to *lem-2Δ* mutant. (E) Left, schematic of reforming NE after first mitotic division in *C. elegans* embryo highlighting endogenous CHMP-7 core enrichment. Right, spinning disk confocal images of GFP^CHMP-7 and mCh:Histone(H)2B at 140 and 400s post anaphase onset. Scale bar, 5 µm (F) Confocal images of fixed 2-cell stage *C. elegans* embryos immunostained with indicated markers (LMN-1 = lamin) in indicated conditions. Yellow arrowheads point to nuclear lobulations. Scale bar, 5 µm. (G) Spinning disk confocal images of NDC-1^mNG and mCh:Histone(H)2B at 600s post anaphase onset in indicated conditions. Scale bar, 5 µm, zoom inset scale bar, 2 µm.

### Barger et al. Figure S3

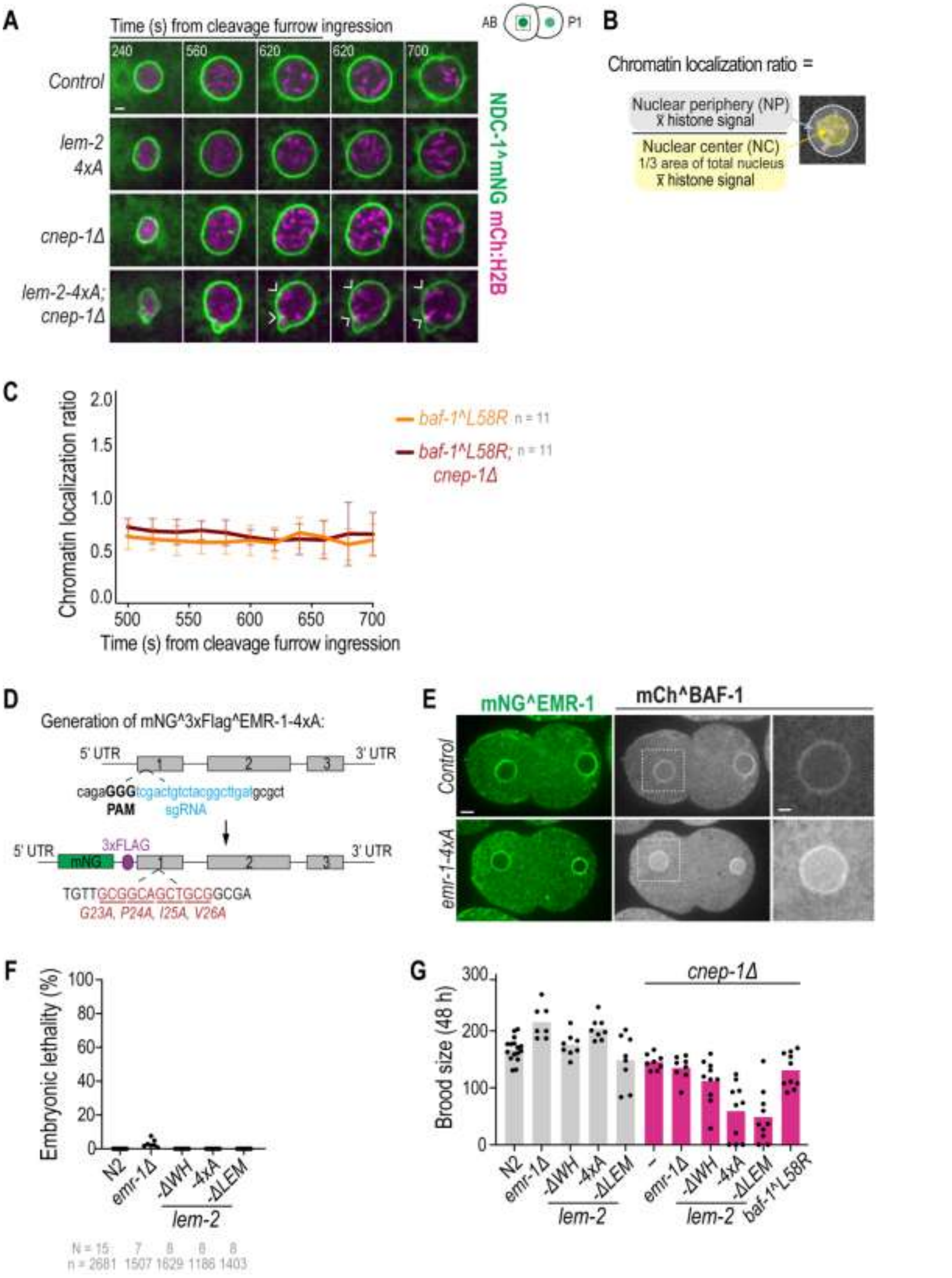

**Figure S3.** (A) Spinning disk confocal images from time lapse series showing NDC-1^mNG and mCh:Histone(H)2B in indicated strains. Time in sec relative to cleavage furrow ingression. White arrowheads point to chromatin localization at the NE relative to Emerin clusters. Scale bar, 2 µm. (B) Schematic of quantification of chromatin localization ratio. (C) Plot (mean ± SD) of chromatin localization ratio in AB nuclei for indicated conditions. Time in sec relative to cleavage furrow ingression (n = # of nuclei). (D) Schematic of CRISPR-Cas9 generation of *emr-1(sbw48[mNG::emr-1-G23A,P24A,I25A,V26A]) I* (E) Spinning disk confocal images of mCh^BAF-1 in mNG^EMR-1 and mNG^EMR-1-4xA mutant. Scale bar, 5 µm, zoom inset scale bar, 2 µm. (F,G) Plot (mean + replicates) of embryonic lethality and brood size in indicated conditions (N = # of worms, n = # of embryos). N, n also shown in Fig. 3I.

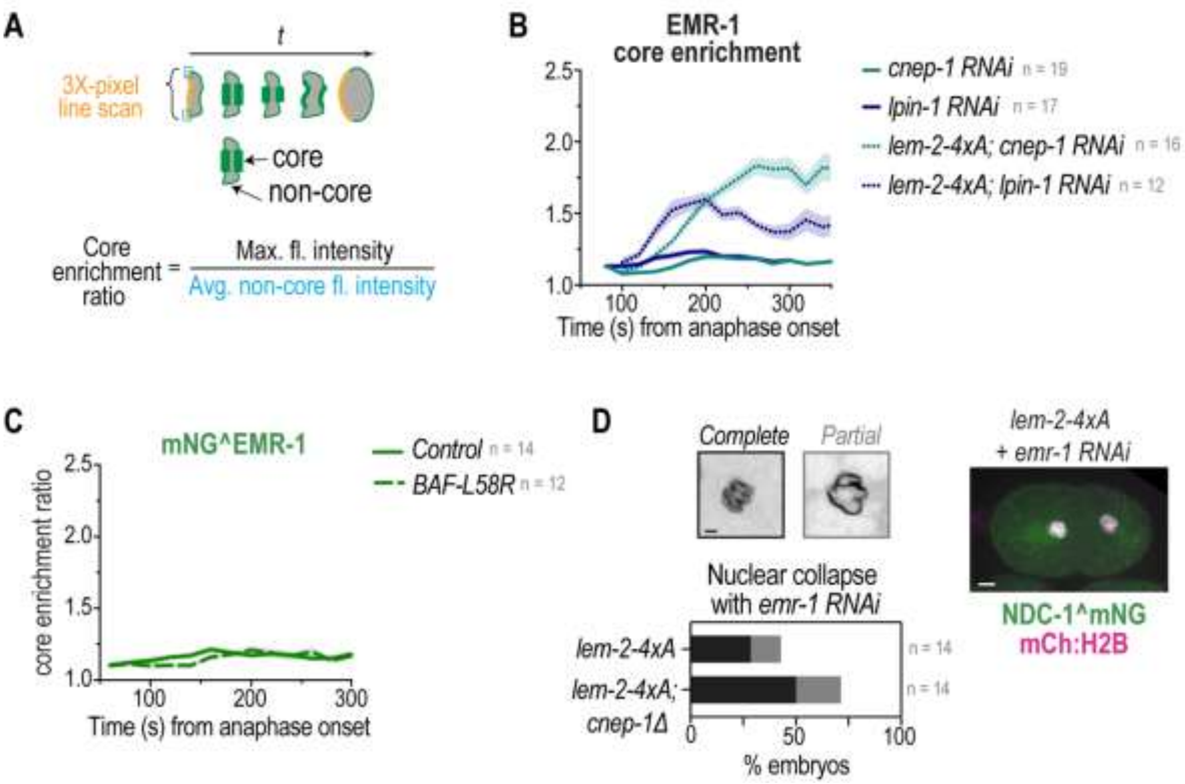


**Figure S4.** (A) Schematic representations of line scan analysis to quantify core and non-core protein dynamics in *C. elegans* embyos (B) Plot (average + SEM) of ratio of maximum fluorescence intensity (core) and non-core fluorescence signal of mNG^EMR-1 measured from anaphase onset (t = 0) in indicated conditions (n = # of embryos). (C) Plot (average + SEM) of ratio of maximum fluorescence intensity (core) and non-core fluorescence signal of mNG^EMR-1 measured from anaphase onset (t = 0) in indicated conditions (n = # of embryos). Control values replicated from Fig. 4C. (D) Left, plot of collapsed nuclear morphology at 600s post anaphase onset in indicated conditions (n = # of embryos). Zoom scale bar, 2 µm. Right, spinning disk confocal images of NDC-1^mNG and mCh:Histone(H)2B at 600s post anaphase onset in indicated condition. Scale bar, 5 µm.

**Barger et al. Figure S5**

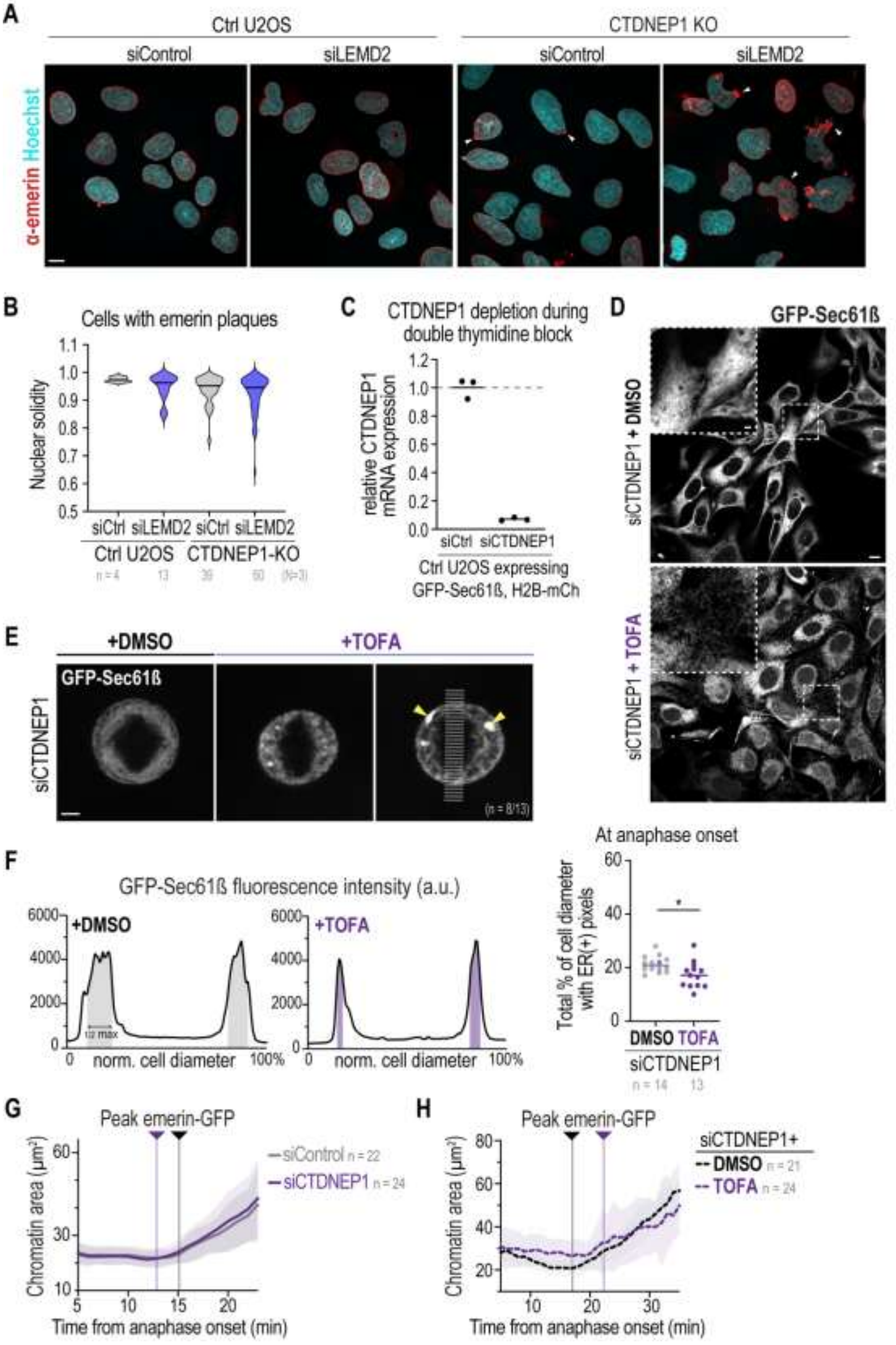

**Figure S5**. (A) Maximum intensity projections of immunostained Control and CTDNEP1-KO U2OS cells with indicated siRNA, immunostained for Emerin. White arrowheads denote Emerin clusters. Scale bar, 10 µm. (B) Violin plot of mean nuclear solidity of cells with emerin clusters in indicated conditions (N = independent experiments, n = cell #). (C) RT-qPCR of CTDNEP1 expression (mean + replicates) in indicated cell line with indicated treatments. Values are normalized to 3B36 expression. Results expressed as fold change in expression relative to siCtrl-treated U2OS values (n = 3 technical replicates). (D) Representative live-cell confocal images of ER morphology in U2OS cells expressing GFP-Sec61β and Histone2B-mCh in indicated conditions. Scale bar, 10 µm, zoom inset scale bar, 2 µm. (E) Representative confocal images of mitotic ER prior to anaphase onset in indicated conditions and placement of 10x pixel line scan. Yellow arrowheads denote ER patches sometimes observed with TOFA-treatment. Scale bar, 5 µm. (F) Left, representative traces of 10x pixel line scan and quantification of mitotic ER at anaphase onset. Right, plot (mean + replicates) of percentage of cell diameter at half maximum value of GFP-Sec61β signal (n = # of cells). * = $p < 0.05$ as determined by Mann-Whitney test. (G, H) Plot (mean ± SD) of chromatin area for indicated conditions in emerin-GFP NE reformation imaging in 5F with timing of peak Emerin-GFP intensity marked (n = nuclei #).

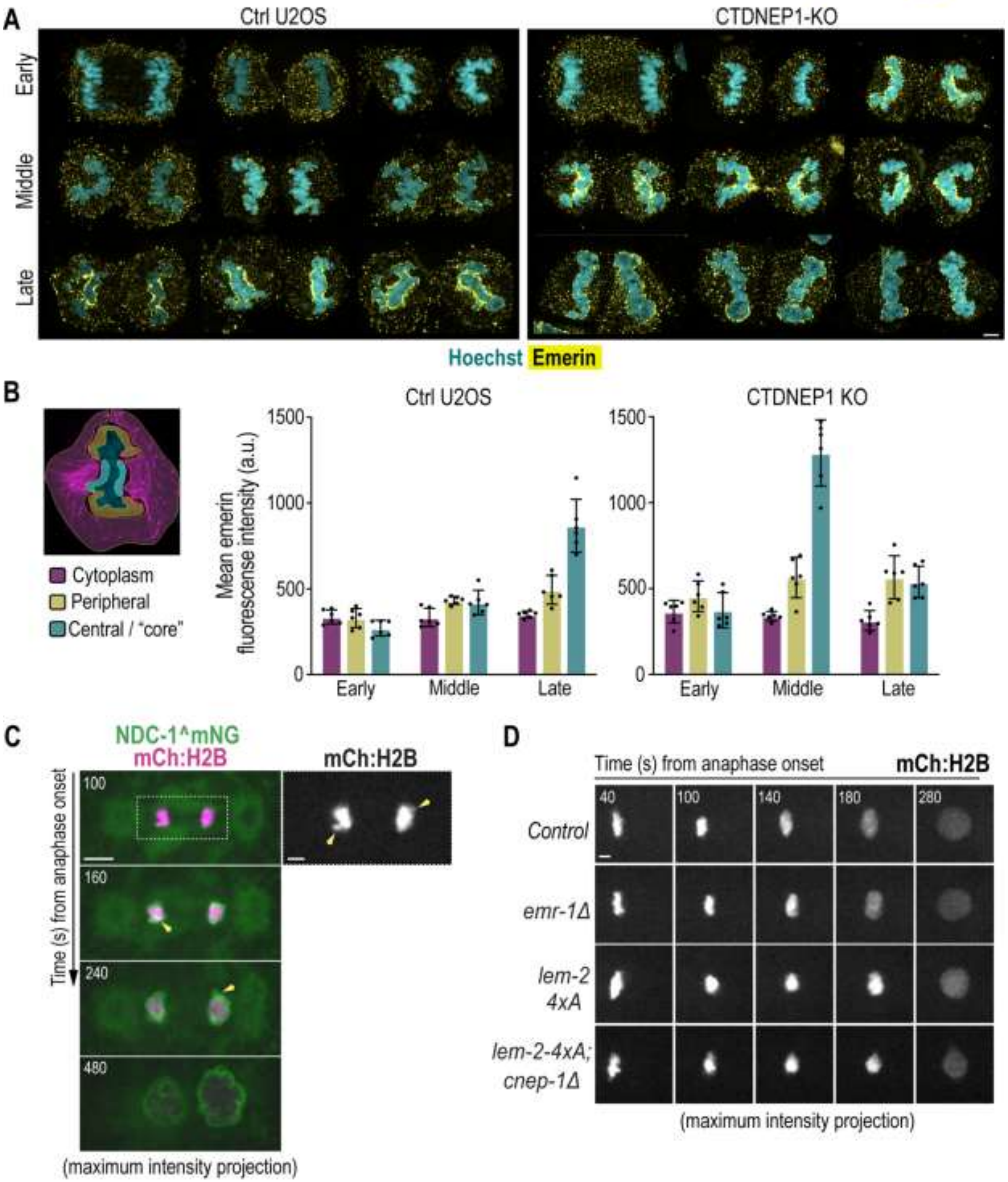


**Figure S6.** (A) Classification of early, middle, late anaphase – telophase in expanded Control and CTDNEP1-KO cells immunostained for Emerin with Hoechst counterstain. Scale bar, 20 µm (not corrected for expansion factor). (B) Left, region of interest to quantify emerin signal at core, non-core and cytoplasm. Right, plot (mean + SD + replicates) of mean emerin intensity in expanded Control and

CTDNEP1-KO cells at indicated locations in time. (C) Spinning disk confocal images from time lapse series showing NDC-1^mNG and mCh:Histone(H)2B in *lem-2-4xA; cnep-1Δ* mutant embryo. Time in sec relative to anaphase onset. Yellow arrowheads point to peripheral chromatin becoming enwrapped. Scale bar, 5 µm. Zoom scale bar, 2 µm. (D) Spinning disk confocal time lapse series showing maximum intensity projection of mCh:Histone(H)2B in indicated strains. Time in sec relative to anaphase onset. Scale bar, 2 µm.

**Barger et al. Figure S7**

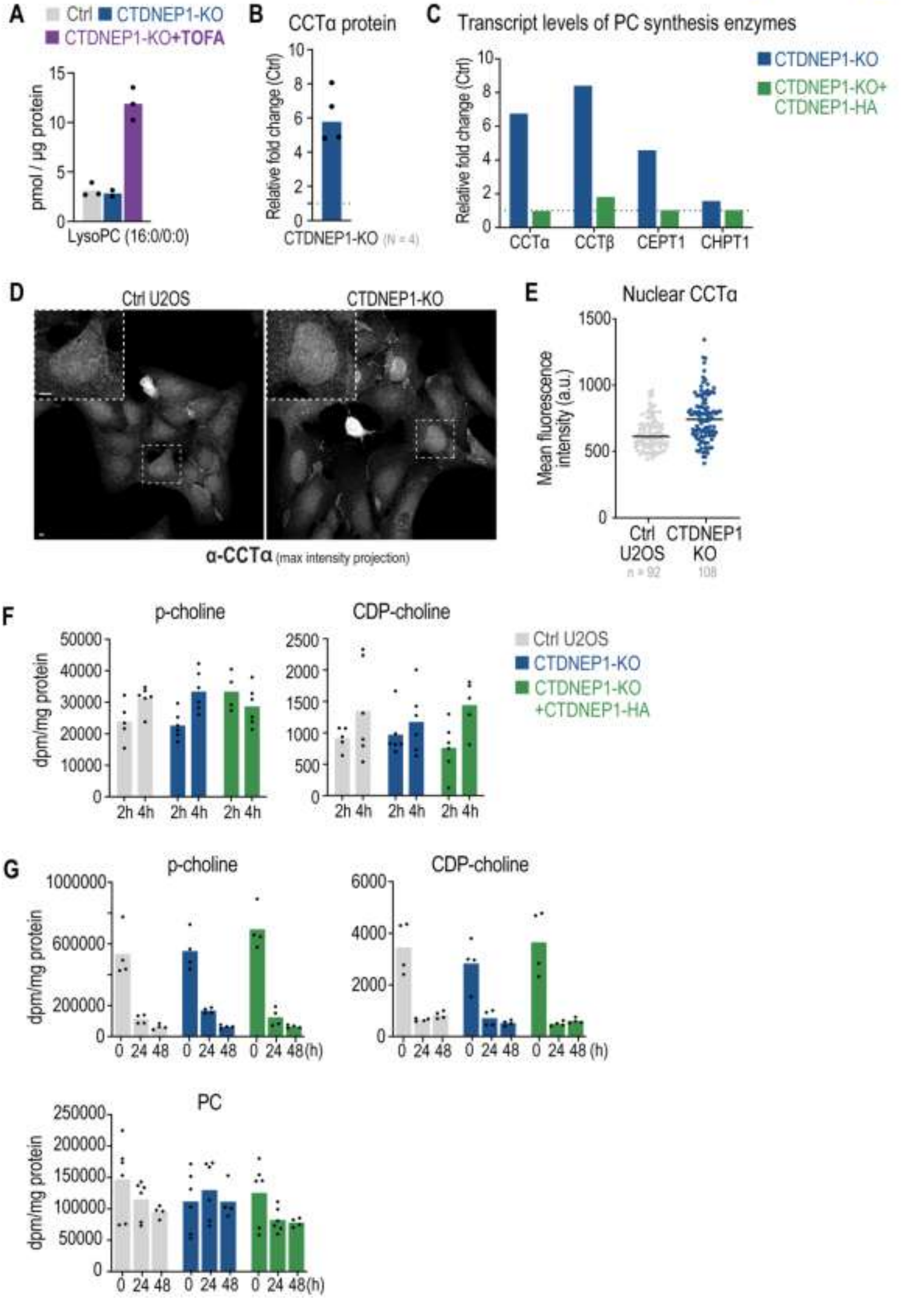

**Figure S7.** (A) Plot (mean + replicates) of lyso-PC in indicated conditions analyzed by lipidomics (n= 2-3 technical replicates). (B) Quantification of CCTα expression by immunoblot (N = 4 biological replicates). (C) qRT-PCR of indicated cells for PC synthesis genes, shown as average fold change in expression relative to mean control U2OS values in 3 technical replicates. (D) Maximum intensity projections of spinning disk confocal images of indicated cells immunostained for CCTα. Scale bar, 10 μm. Zoom inset scale bar, 5 μm. (E) Plot (mean + replicates) of mean fluorescence intensity quantification of nuclear CCTα. Showing representative data from 3 independent experiments (n = cell #). (F) Plot (mean + replicates) of [$^{3}$H]Choline incorporation into PC metabolites for indicated cells at indicated time points (N = 4-6 biological replicates). (G) Plot (mean + replicates) of [$^{3}$H]Choline incorporation into PC and metabolites for indicated cells at indicated time points (N = 4-6 biological replicates).

#### Supplemental Movie Legends

**Movie S1. Formation of nuclear lobulations during NE assembly in *C. elegans* embryos**, related to Figure 2F. Confocal time series of nuclear assembly in first cell division of *C. elegans* embryos. Endogenous NDC-1^mNG (green) marks the nuclear envelope and mCherry:Histone2B (magenta) labels chromatin. Images were acquired every 20 s – playback rate is 80X real time. Scale bar, 2 μm.

**Movie S2. Membrane invasions are fenestrated and adjacent to membrane stacks in *lem-2-4xA*; *cnep-1Δ* double mutant embryos**, related to Figure 3B. 3D electron tomography of nuclear membrane invasion in *lem-2-4xA*; *cnep-1Δ* double mutant embryos with nuclear membranes traced (green). Scale bar in tomography, 600 nm, scale bar in model tracing 200 nm.

**Movie S3. Emerin clustering and chromatin tethering in *C. elegans* embryos**, related to Figure 3F. Confocal time series of nuclei at 2-cell stage of *C. elegans* embryos expressing endogenous mNG^EMR-1 (green) and mCherry:Histone2B (magenta) in indicated strains. Emerin clusters are indicated by white arrowheads. Images were acquired every 20 s. Scale bar, 2 μm.

**Movie S4. Core domain localization of LEM-2^mNG and mNG^EMR-1 in *C. elegans* first cell division**, related to Figure 4A. Confocal time series of *C. elegans* embryos expressing either endogenous LEM-2^mNG or mNG^EMR-1. Images were acquired every 20 s – playback rate is 80X real time. Scale bar, 5 μm.

**Movie S5. Emerin dynamics during NE assembly in siControl and siCTDNEP1 cells**, related to Figure 5E-F. Confocal time series of U2OS cells stably-expressing emerin-GFP (gray) stained with 5'-SiR-Hoechst (blue) and treated with control and siCTDNEP1 siRNA. Images were acquired every minute – playback rate is 249X real time. Scale bar, 5 μm.

**Movie S6. ER membranes infiltrate chromatin mass during NE assembly with depletion of CTDNEP1**, related to Figure S6A. Confocal time series of U2OS cells stably-expressing GFP-Sec61β (green) to label ER and H2B-mCherry (magenta) to label chromatin, treated with control and siCTDNEP1 siRNA. Images were acquired every 30 sec – playback rate is 123X real time. Scale bar, 5 μm.

**Movie S7. TOFA treatment reduces ER membrane infiltration into mitotic chromosomes during NE reformation,** related to Figure 6A. Confocal time series of U2OS cells stably-expressing GFP-Sec61β (green) to label ER and H2B-mCherry (magenta) to label chromatin, treated siCTDNEP1 siRNA and DMSO or TOFA. Images were acquired every 30 sec – playback rate is 115X real time. Scale bar, 5 µm.

**Movie S8. Expansion microscopy of control and CTDNEP1-KO cells at maximal emerin enrichment**, related to Figure 6C. Representative ExM image of control and CTDNEP1-KO cell stained for emerin (yellow), tubulin (magenta) and DNA (cyan) during NE reformation. Scale bar, 20 µm.

**Movie S9. Expansion microscopy of mitotic chromatin showcasing emerin localization**, related to Figure 6F. Chromatin mass of daughter nuclei in late anaphase in control and CTDNEP1-KO cells with emerin (yellow) and Hoechst (cyan) staining. Scale bar, 20 µm.

Table S1. *C. elegans* strains used in this study

| ***C. elegans* strain name** | **Genotype** | **Source** |
|---|---|---|
| N2 | (Bristol): Wild-type (ancestral) | Caenorhabditis Genetics Center (CGC) |
| SBW448 | lem-2(sbw20) II | Barger et al., 2023 |
| SBW317 | lem-2(sbw17[G24A, P25A, V26A, T27A]) II | This study |
| SBW318 | lem-2(syb4612) II | This study; Suny Biotech |
| SBW319 | lem-2(syb4718) II | Barger et al., 2023 |
| SBW466 | lem-2(sbw20) II; ltIs24 [pAZ132; pie-1p::GFP::tba-2; unc-119 (+)]; unc-119(ed3)III; sbwSi16 [pSB486; lem-2 5'UTR::mScarlet-i-lem-2::lem-2 3'UTR] cb-unc-119(+) IV | This study |
| SBW467 | lem-2(sbw20) II; ltIs24 [pAZ132; pie-1p::GFP::tba-2; unc-119 (+)]; unc-119(ed3)III; sbwSi19 [pSB694; lem-2 5'UTR::mScarlet-i-lem-2(G24A, P25A, V26A, T27A)::lem-2 3'UTR] cb-unc-119(+) IV. | This study |
| SBW468 | lem-2(sbw20) II; ltIs24 [pAZ132; pie-1p::GFP::tba-2; unc-119 (+)]; unc-119(ed3)III; sbwSi17 [pSB488; lem-2 5'UTR::mScarlet-i-lem-2-ΔLEM::lem-2 3'UTR] cb-unc-119(+) IV. | This study |
| SBW469 | lem-2(sbw20) II; ltIs24 [pAZ132; pie-1p::GFP::tba-2; unc-119 (+)]; unc-119(ed3)III; sbwSi15 [pSB487; lem-2 5'UTR::mScarlet-i-lem-2-ΔWH::lem-2 3'UTR] cb-unc-119(+) IV | This study |
| SBW414 | gfp::chmp-7 II; unc-119(ed3) III; ltIs37 [pAA64 pie-1p::mCherry::his-58; unc-119 (+)] IV | Barger et al., 2023 |
| SBW417 | gfp::chmp-7 II; lem-2(sbw17[G24A, P25A, V26A, T27A]) II; unc-119(ed3) III; ltIs37 [pAA64; pie-1p::mCherry::his-58; unc-119 (+)] IV | This study |
| SBW418 | gfp::chmp-7 II; lem-2(syb4612) II; unc-119(ed3) III; ltIs37 [pAA64; pie-1p::mCherry::his-58; unc-119 (+)] IV | This study |

| | | |
|---|---|---|
| SBW419 | gfp::chmp-7 II; lem-2 (syb4718) II; unc-119(ed3) III; ltIs37 [pAA64; pie-1p::mCherry::his-58; unc-119 (+)] IV | Barger et al., 2023 |
| SBW313 | unc-119(ed3) III; ltIs37 [pAA64; pie-1p::mCherry::his-58; unc-119 (+)] IV; ndc-1(sbw15[ndc1::mNG]) V | This study |
| SBW429 | emr-1(gk119) I; unc-119(ed3) III; ltIs37 [pAA64; pie-1p::mCherry::his-58; unc-119 (+)] IV; ndc-1(sbw15[ndc1::mNG]) V | This study |
| SBW456 | lem-2(sbw20) II; unc-119(ed3) III; ltIs37 [pAA64; pie-1p::mCherry::his-58; unc-119 (+)] IV; ndc-1(sbw15[ndc1::mNG]) V | This study |
| SBW434 | baf-1(syb2609[L58R]) III; unc-119(ed3) III; ltIs37 [pAA64; pie-1p::mCherry::his-58; unc-119 (+)] IV; ndc-1(sbw15[ndc1::mNG]) V | This study |
| SBW360 | lem-2(sbw17[G24A, P25A, V26A, T27A]) II; unc-119(ed3) III; ltIs37 [pAA64; pie-1p::mCherry::his-58; unc-119 (+)] IV; ndc-1(sbw15[ndc1::mNG]) V | This study |
| SBW362 | lem-2(syb4718) II; unc-119(ed3) III; ltIs37 [pAA64; pie-1p::mCherry::his-58; unc-119 (+)] IV; ndc-1(sbw15[ndc1::mNG]) V | This study |
| SBW364 | scpl-2(tm4369) II; unc-119(ed3) III; ltIs37 [pAA64; pie-1p::mCherry::his-58; unc-119 (+)] IV; ndc-1(sbw15[ndc1::mNG]) V | This study |
| SBW655 | emr-1(gk119) I; scpl-2(tm4369) II; unc-119(ed3) III; ltIs37 [pAA64; pie-1p::mCherry::his-58; unc-119 (+)] IV; ndc-1(sbw15[ndc1::mNG]) V | This study |
| SBW435 | scpl-2(tm4369) II; baf-1(syb2609[L58R]) III; unc-119(ed3) III; ltIs37 [pAA64; pie-1p::mCherry::his-58; unc-119 (+)] IV; ndc-1(sbw15[ndc1::mNG]) V | This study |
| SBW411 | lem-2(sbw17[G24A, P25A, V26A, T27A]) II; scpl-2(tm4369) II; unc-119(ed3) III; ltIs37 [pAA64; pie-1p::mCherry::his-58; unc-119 (+)] IV; ndc-1(sbw15[ndc1::mNG]) V | This study |

| | | |
|---|---|---|
| SBW413 | lem-2(syb4718) II; scpl-2(tm4369) II; unc-119(ed3) III; ltIs37 [pAA64; pie-1p::mCherry::his-58; unc-119 (+)] IV; ndc-1(sbw15[ndc1::mNG]) V | This study |
| SBW506 | cnep-1(syb6829 [D63E, D65T]) II; unc-119(ed3) III; ltIs37[pAA64; pie-1p::mCherry::his-58; unc-119 (+)] IV; ndc-1(sbw15[ndc1::mNG]) V | This study |
| SBW507 | lem-2(sbw17[G24A, P25A, V26A, T27A]) II; cnep-1(syb6829 [D63E, D65T]) II; unc-119(ed3) III; ltIs37[pAA64; pie-1p::mCherry::his-58; unc-119 (+)] IV; ndc-1(sbw15[ndc1::mNG]) V | This study |
| SBW622 | lem-2(sbw17[G24A, P25A, V26A, T27A]) II; scpl-2(tm4369) II; mel-28(bq5[GFP::mel-28]) III ; unc-119(ed3) III; ltIs37 [pAA64; pie-1p::mCherry::his-58; unc-119 (+)] IV | This study |
| SBW256 | scpl-2(tm4369) II | Bahmanyar et al., 2014 |
| SBW32 | ltIs24 [pAZ132; pie-1p::GFP::tba-2; unc-119 (+)]; ltIs37 [pAA64; pie-1p::mCherry::his-58; unc-119 (+)] IV | (Penfield et al., 2018) |
| SBW49 | scpl-2(tm4369) II ; ltIs24 [pAZ132; pie-1p::GFP::tba-2; unc-119 (+)]; ltIs37 [pAA64; pie-1p::mCherry::his-58; unc-119 (+)] IV | (Penfield et al., 2020) |
| SBW455 | lem-2(sbw20) II; ltIs24 [pAZ132; pie-1p::GFP::tba-2; unc-119 (+)]; ltIs37 [pAA64; pie-1p::mCherry::his-58; unc-119 (+)] IV | Barger et al., 2023 |
| SBW63 | chmp-7(hz12) II; ltIs24 [pAZ132; pie-1p::GFP::tba-2; unc-119 (+)]; ltIs37 [pAA64; pie-1p::mCherry::his-58; unc-119 (+)] IV | (Penfield et al., 2020) |
| SBW480 | emr-1(gk119) I; ltIs24 [pAZ132; pie-1p::GFP::tba-2; unc-119 (+)]; ltIs37 [pAA64; pie-1p::mCherry::his-58; unc-119 (+)] IV | Barger et al., 2023 |
| SBW238 | ltIs24 [pAZ132; pie-1p::GFP::tba-2; unc-119 (+)]; ltIs37 [pAA64; pie-1p::mCherry::his-58; unc-119 (+)] IV; spas-1(tm638) V | This study |
| SBW461 | lem-2(sbw20) II; scpl-2(tm4369) II ; ltIs24 [pAZ132; pie-1p::GFP::tba-2; unc-119 (+)]; ltIs37 [pAA64; pie-1p::mCherry::his-58; unc-119 (+)] IV | This study |

| | | |
|---|---|---|
| SBW75 | chmp-7(hz12) II; scpl-2(tm4369) II; ltIs24 [pAZ132; pie-1p::GFP::tba-2; unc-119 (+)]; ltIs37 [pAA64; pie-1p::mCherry::his-58; unc-119 (+)] IV | (Penfield et al., 2020) |
| SBW477 | emr-1(gk119) I; scpl-2(tm4369) II; ltIs24 [pAZ132; pie-1p::GFP::tba-2; unc-119 (+)]; ltIs37 [pAA64; pie-1p::mCherry::his-58; unc-119 (+)] IV | This study |
| SBW664 | scpl-2(tm4369) II; unc-119(ed3) III; ltIs37 [pAA64; pie-1p::mCherry::his-58; unc-119 (+)] IV; ndc-1(sbw15[ndc1::mNG]) V; spas-1(tm638) V | This study |
| SBW334 | lem-2(sbw17[G24A, P25A, V26A, T27A]) II; ltIs24 [pAZ132; pie-1p::GFP::tba-2; unc-119 (+)]; ltIs37 [pAA64; pie-1p::mCherry::his-58; unc-119 (+)] IV | This study |
| SBW338 | lem-2(syb4718) II; ltIs24 [pAZ132; pie-1p::GFP::tba-2; unc-119 (+)]; ltIs37 [pAA64; pie-1p::mCherry::his-58; unc-119 (+)] IV | This study |
| SBW356 | lem-2(sbw17[G24A, P25A, V26A, T27A]) II; scpl-2(tm4369) II; ltIs24 [pAZ132; pie-1p::GFP::tba-2; unc-119 (+)]; ltIs37 [pAA64; pie-1p::mCherry::his-58; unc-119 (+)] IV | This study |
| SBW358 | lem-2(syb4718) II; scpl-2(tm4369) II; ltIs24 [pAZ132; pie-1p::GFP::tba-2; unc-119 (+)]; ltIs37 [pAA64; pie-1p::mCherry::his-58; unc-119 (+)] IV | This study |
| SBW459 | lem-2(sbw20) II; scpl-2(tm4369) II | This study |
| SBW400 | lem-2(sbw17[G24A, P25A, V26A, T27A]) II; scpl-2(tm4369) II | This study |
| SBW401 | lem-2(syb4612) II; scpl-2(tm4369) II | This study |
| SBW402 | lem-2 (syb4718) II; scpl-2(tm4369) II | This study |
| SBW169 | scpl-2(tm4369) II; baf-1(syb2609[L58R]) III | This study |
| SBW312 | emr-1(sbw16[mNG::3xFlag::emr-1]) I; ltIs37 [pAA64; pie-1p::mCherry::his-58; unc-119 (+)] IV | Barger et al., 2023 |
| SBW146 | lem-2(sbw5)[lem-2::mNG]) II; unc-119(ed3) III; ltIs37 [pAA64; pie-1p::mCherry::his-58; unc-119 (+)] IV | Barger et al., 2023 |
| SBW754 | emr-1(sbw16[mNG::3xFlag::emr-1]) I; lem-2(sbw17[G24A, P25A, V26A, T27A]) II; ltIs37 [pAA64; pie-1p::mCherry::his-58; unc-119 (+)] IV | This study |

| | | |
|---|---|---|
| SBW350 | emr-1(sbw16[mNG::3xFlag::emr-1]) I; baf-1(syb2609[L58R]) III; ltIs37 [pAA64; pie-1p::mCherry::his-58; unc-119 (+)] IV | Barger et al., 2023 |
| SBW795 | emr-1(sbw16[mNG::3xFlag::emr-1]) I; baf-1(bq13[mCherry:baf-1]) III | This study |
| SBW796 | emr-1(sbw48[mNG::emr-1-G23A,P24A,I25A,V26A]) I; baf-1(bq13[mCherry:baf-1]) III | This study |

Table S2. Oligonucleotides used in this study.

| **Oligonucleotides for dsRNA production** | | | |
|---|---|---|---|
| Gene | Oligonucleotide 1 | Oligonucleotide 1 | Template |
| H37A05.1 (*lpin-1*) | AATTAACCCTCACTAAAGGG GCCATTTGTCCACTCTCAT | TAATACGACTCACTATAGGCT TACACACTCGGCGGTTTT | N2 genomic |
| F45E12.1 (*cnep-1*) | AATTAACCCTCACTAAAGGAT GACAACAATCGCACAGTCT | TAATACGACTCACTATAGGGA GACGGAGAAGGTGTAGAAC | cDNA |
| M01D7.6 (*emr-1*) | AATTAACCCTCACTAAAGGCG A ACTACGCGATAGCCTTA | TAATACGACTCACTATAG GCCCAAGAATCCTCCTTT GTT | N2 genomic |
| **Oligonucleotides for RT-qPCR** | | | |
| Reagent | Oligonucleotide 1 | Oligonucleotide 1 | Source |
| Hs CTDNEP1 qPCR primer | CATTTACCTTCTGCGGAG GC | CACCTGGGCTAGCCGATT C | Merta, Carasquillo Rodriguez et al., 2021). |
| Hs 36B4 qPCR primer | AACATGCTCAACATCTCCCC | CCGACTCCTCCGACTCTTC | Neuhaus et al., 2011 |
| Hs PCYT1A qPCR primer | GCCCTATGTCAGGGTAACTAT GG | GCGTGACCAGAGTGAAATAA GT | Dorighello et al., 2022 |
| Hs PCYT1B qPCR primer | ATGCAAGAGCCCTTATGCAAG | GCTTCGTATCTCTCGGCTTCA TT | Dorighello et al., 2022 |

| | | | |
|---|---|---|---|
| Hs CEPT1 qPCR primer | ATGTGGAGATTCTCACCCGGA | TCTTCTAGCCGCTTTAGTTGGT | Dorighello et al., 2022 |
| Hs CHPT1 qPCR primer | TTGCCGCTCGCTTAGGAAC | GCCTGAAACATAAGTCTGCCAAT | Dorighello et al., 2022 |

Table S3. Recombinant DNA or ssODN used in this study.

| **Identifier** | **Recombinant DNA** | **Source** |
|---|---|---|
| pSB486 | pCFJ151-lem-2-5'UTR-mScarlet-i-LEM-2-lem-2 3'UTR | This study |
| pSB694 | pCFJ151-lem-2-5'UTR-mScarlet-i-lem-2(G24A, P25A, V26A, T27A)-lem-2 3'UTR | This study |
| pSB487 | pCFJ151-lem-2-5'UTR-mScarlet-i-LEM-2-ΔWH-lem-2 3'UTR | This study |
| pSB488 | pCFJ151-lem-2-5'UTR-mScarlet-i-LEM-2-ΔLEM -lem-2 3'UTR | This study |
| pSB514 | pDD122-emr-1 | Barger et al., 2023 |
| pSB961 | pDD268-emr-1(G23A, P24A, I25A, V26A]) | This study |
| *lem-2* crRNA | GGTAACTGGAACGACTCGTT | IDT |
| *lem-2-4xA* repair template | GCCGAGTTGAACGTTCGCGGGGCGAATGTTGCGGCCGCAGCTGGAACGACTCGTTCGCTGTACGAGAAGAAG | Keck Oligos |
